# Physics-informed deep generative learning for quantitative assessment of the retina

**DOI:** 10.1101/2023.07.10.548427

**Authors:** Emmeline Brown, Andrew Guy, Natalie Holroyd, Paul Sweeney, Lucie Gourmet, Hannah Coleman, Claire Walsh, Athina Markaki, Rebecca Shipley, Ranjan Rajendram, Simon Walker-Samuel

**Affiliations:** Centre for Computational Medicine, University College London, London, UK; Moorfields Eye Hospital, London, UK; Department of Engineering, University of Cambridge, Cambridge, UK; Cancer Research UK Cambridge Institute, University of Cambridge, Cambridge, UK; Department of Mechanical Engineering, University College London, London, UK; Institute of Ophthalmology, University College London, UK

## Abstract

Disruption of retinal vasculature is linked to various diseases, including diabetic retinopathy and macular degeneration, leading to vision loss. We present here a novel algorithmic approach that generates highly realistic digital models of human retinal blood vessels based on established biophysical principles, including fully-connected arterial and venous trees with a single inlet and outlet. This approach, using physics-informed generative adversarial networks (PI-GAN), enables the segmentation and reconstruction of blood vessel networks that requires no human input and out-performs human labelling. Our findings highlight the potential of PI-GAN for accurate retinal vasculature characterization, with implications for improving early disease detection, monitoring disease progression, and improving patient care.

## Introduction

Disruption of retinal vasculature is associated with a range of diseases which can result in loss of vision, including diabetic retinopathy (DR) [1] and macular degeneration [2]. It is also increasingly recognized that retinal vasculature can indicate the presence of systemic pathology, such as vascular dementia [3] and cardiovascular disease [4]. Automated methods to characterize changes in retinal vasculature from clinical imaging data therefore offer substantial promise for high-throughput, early detection of disease [5], which is critically required to meet the increasing incidence of retinal disease, potentially alongside other vascular diseases, and their associated burden on healthcare systems [6].

Much attention has been placed on supervised deep learning in this regard, where deep neural networks are trained to categorise images according to diagnosis or identify the location of features of interest [7]. Supervised learning, particularly with U-net architectures [8], first rose to prominence in retinal image analysis for segmenting retinal layers in optical coherence tomography (OCT) data [9], alongside blood vessels segmentation in retinal photographs [10, 11]. A significant limitation to this type of approach is the lack of high-quality, manually-labelled image data in sufficient quantities to enable accurate and generalisable predictions to be made [12]. This problem is particularly acute for the detection of blood vessels, in which manual labelling is highly time-consuming, generally limited to two-dimensional (2D) projections, confined to larger vessels only, and generally does not distinguish between arteries and veins [13].

To address these challenges, we describe here a novel set of algorithms that can generate highly realistic digital models of human retinal blood vessels, using established biophysical principles and unsupervised deep learning. Our biophysical models capture the complex structure of retinal vasculature, with interconnecting arterial and venous trees that are inherently three-dimensional, multi-scale and fully inter-connected via a capillary bed. They also feature dedicated macula and optic disc features. The central biophysical principles we draw on are 1) Murray’s Law, in which vessel diameters, branching distances and branching angles are optimised to form a balance between pumping power and blood volume and minimize resistance to flow [14]; and 2) fluid dynamics to model blood flow and vascular exchange. The latter is made possible by our synthetic networks containing fully-connected arterial and venous trees with a single inlet and outlet (the central retinal artery and vein), allowing blood flow and contrast agent delivery (e.g. fluorescein) data to be simulated with minimal assumptions in regard to network boundary conditions.

In this work, we investigate whether, through the use of generative deep learning, our biophysics-informed vascular network models can be used to infer information from real-world retinal images, such as the segmentation and reconstruction of blood vessel networks, without the need to perform any manual labelling, in an approach termed physics-informed generative adversarial learning (PI-GAN) [15]. An overview of our framework is provided in **Figure 1**. Generative adversarial networks (GANs) incorporating cycle-consistency have previously been used for medical imaging domain machine learning tasks such as chest MRI to X-ray CT transformation [16], PET image denoising [17], and artefact reduction in fundus photography [18]. Likewise Menten et al used the space colonisation algorithm to generate macular blood vessel images, which they coupled with deep learning [19].

**Figure 1.**
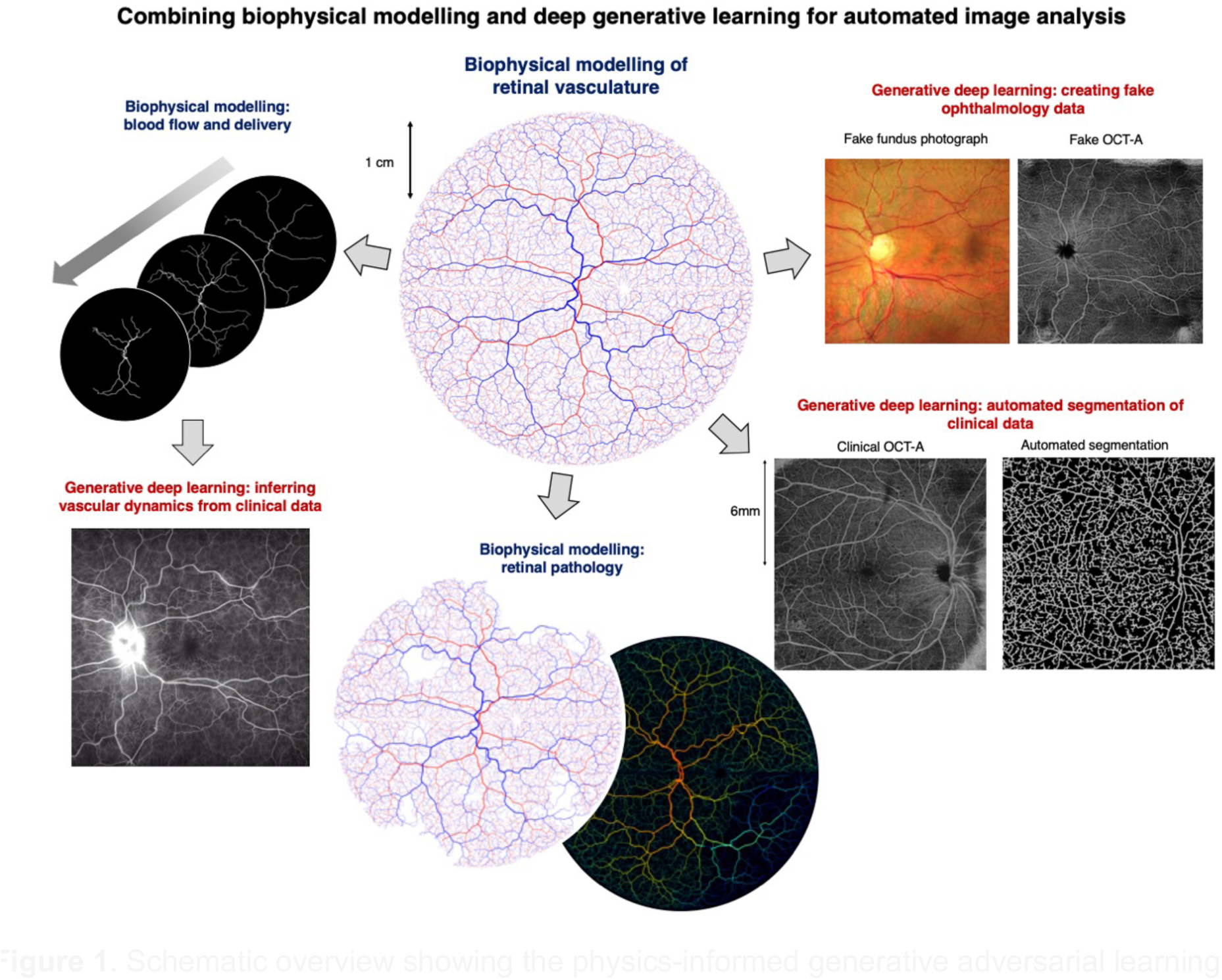
Schematic overview showing the physics-informed generative adversarial learning (PI-GAN) framework developed in this study. Retinal blood vessel networks, featuring arterial and venous trees connected by a capillary bed, and special treatment of macular and optic disc features, were simulated using space filling growth algorithms based on Murray’s law. Blood flow and fluorescein delivery were simulated in synthetic vascular networks, using one-dimensional Poiseuille flow. By combining this with cycle-consistent, physics-informed deep generative learning, vessel simulations were converted into synthetic medical image data (fundus photography, Optical coherence tomography angiography (OCT-A) and fluorescein angiography), and the same trained networks used to detect blood vessels in clinical images.

We demonstrate here the ability of our retinal simulation framework to accurately simulate real-world retinal vasculature, including blood flow, and model the presentation of two common vascular pathologies: DR and retinal vein occlusion (RVO). Moreover, we show that our use PI-GAN workflow allows retinal vasculature to be segmented without any human manual labelling, and which outperforms state-of-the-art supervised learning approaches. This therefore offers numerous opportunities for improved detection and quantification of retinal disease in clinical ophthalmology.

## Results

### Procedural modelling of retinal vasculature

Retinal vascular networks were simulated in multiple, linked steps, using a combination of algorithms that draw on the known geometry and biophysics of retinal vasculature. In total, our procedural model of retinal vasculature contained 26 parameters (**Supplemental Table 1**), each of which were randomly sampled to simulate the broad range of retinal geometries occurring in the population (**Figure 2a-c**) [20, 21].

**Figure 2:**
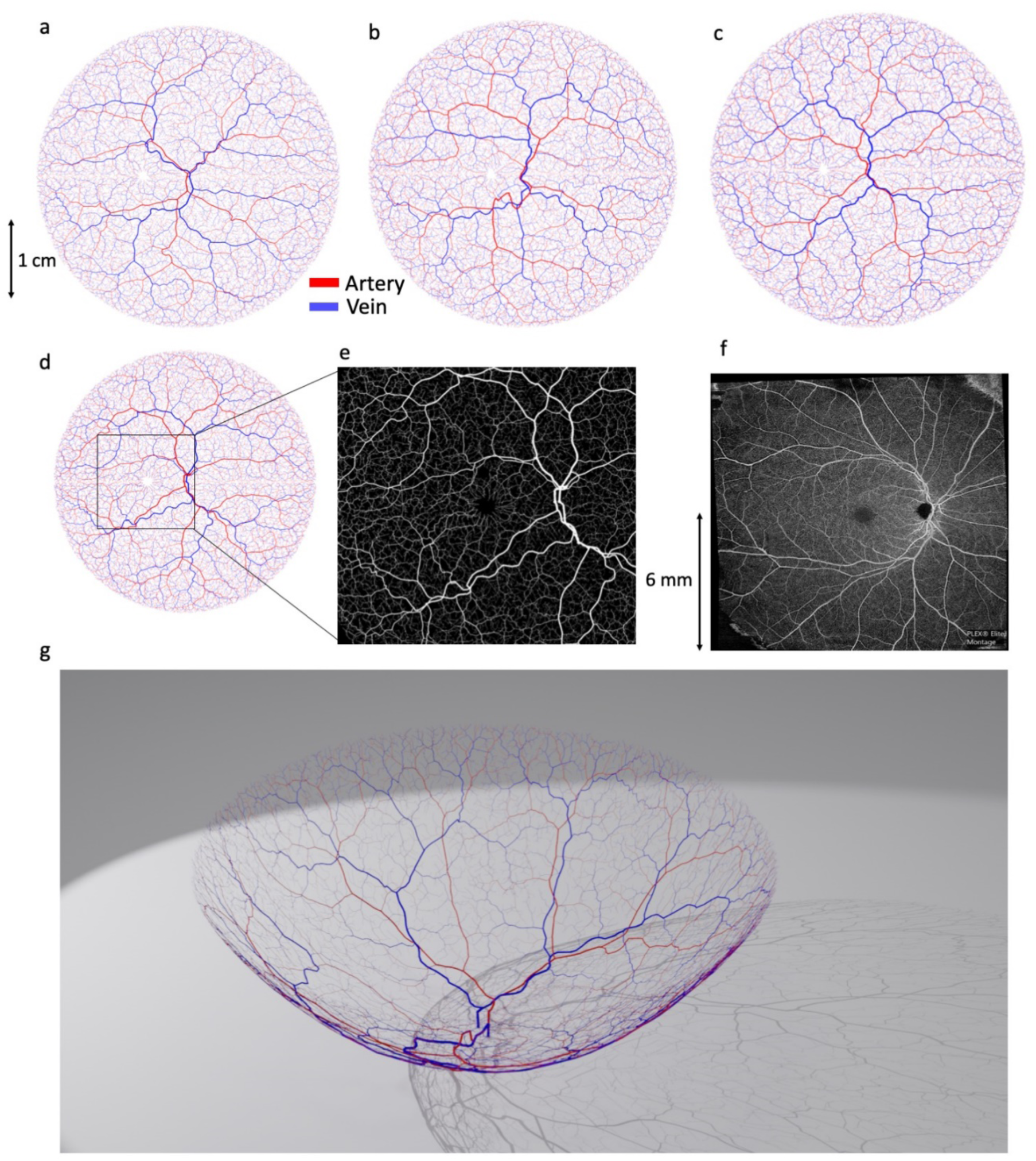
Procedural generation of retinal vasculature using constrained constructive optimisation and lattice sequence vascularisation. a-c) Examples of synthetic retinal vascular networks, featuring arterial (red) and venous (blue) trees, and with geometry optimised according to Murray’s law. Each simulation run used a different set of physiological parameter values, randomly sampled from the distributions defined in Supplemental Table 1. d-f) A synthetic retina (d) with a 12×12 mm region surrounding the optic disc and macula (e) compared with a real OCT-A image (f). g) A simulated vascular network projected onto three-dimensional surface.

Networks were seeded using a Lindenmayer-system (L-system) [22], in which initial central retinal artery and vein segments were positioned at the location of an optic disc and iteratively branched within a plane. The first arterial and venous segment radii were 135 ± 15 µm and 151 ± 15 µm, respectively [23]. Branching was performed asymmetrically to create characteristically large vessels surrounding the macula, with smaller branches reaching towards the periphery, as observed in retinal images [23].

Seeding L-systems were then grown to the edge of the retina using a variant of constrained constructive optimisation (CCO) [24–28]. This step transformed L-system networks into realistic, space-filling networks with geometries defined by Murrays law [14] (exponent of 2.4 ± 0.11 [29]), whilst retaining the realistic macroscopic branching geometry imposed by the L-system seeding (**Figure 2d-f**). A final growth step was incorporated to create the characteristic branching pattern of the macula, with radial alignment of arterioles and venules, greater relative vascular flow density (between 1.5 and 2.0 times the perfusion fraction) and a central avascular fovea.

Following growth, we augmented vessels with sinusoidal displacements to replicate the tortuous vasculature commonly observed in human retinas, with a greater displacement imposed on veins. A continuous capillary bed was generated using either 1) a 2D Voronoi algorithm that arterial and venous endpoints were connected to [30] or 2) a 2D space colonization algorithm [31]. Following simulation within a 2D plane, vessels were projected onto a hemispherical mesh (radius 23 - 25 mm) featuring macula and optic disc structures generated using a mixed Gaussian profile [32] (**Figure 2, Supplemental Figure 1**).

### Comparison of synthetic networks with real-world networks

Our set of retinal network growth algorithms is designed to provide an authentic replication of real retinal vasculature, by following established biophysical principles. To quantitatively evaluate the accuracy of these synthetic networks, we manually labelled all visible blood vessels in 19 optical coherence tomography angiography (OCT-A) image datasets, using in-house software. This included differentiating arteries and veins (A-V) in a subset of images (n=5), using retinal photographs as a reference for determining A-V status. Vessel branching angle, inter-branch length, tortuosity, and radius were measured in three regions: the macula, the vessels surrounding the optic disc, and the periphery. The macula was defined as a 5.5 mm diameter circular area centred on the fovea, based on measurements referenced in Remington and Goodwin [33]. The vessels surrounding the optic disc were labelled as a 3.6mm diameter centred at the optic disc, due to mean vertical and horizontal diameters of the optic disc reported as 1.88 and 1.77mm respectively [34]. Vessels outside these regions were defined as ‘peripheral’. 100 synthetic retinal networks were initially created, with parameter values randomly drawn from the ranges shown in **Supplemental Table 1**.

According to ANOVA analysis, all geometric parameters associated with synthetic blood vessel networks did not reach the level of statistical significance compared to those measured in normal controls using manual segmentation of OCT-A images (branching angle, p = 0.82; vessel length, p = 0.17; vessel tortuosity, p = 0.095; vessel network volume, p = 0.061; vessel diameter, p = 0.59) (**Figure 3, Supplemental Table 2**).

**Figure 3.**
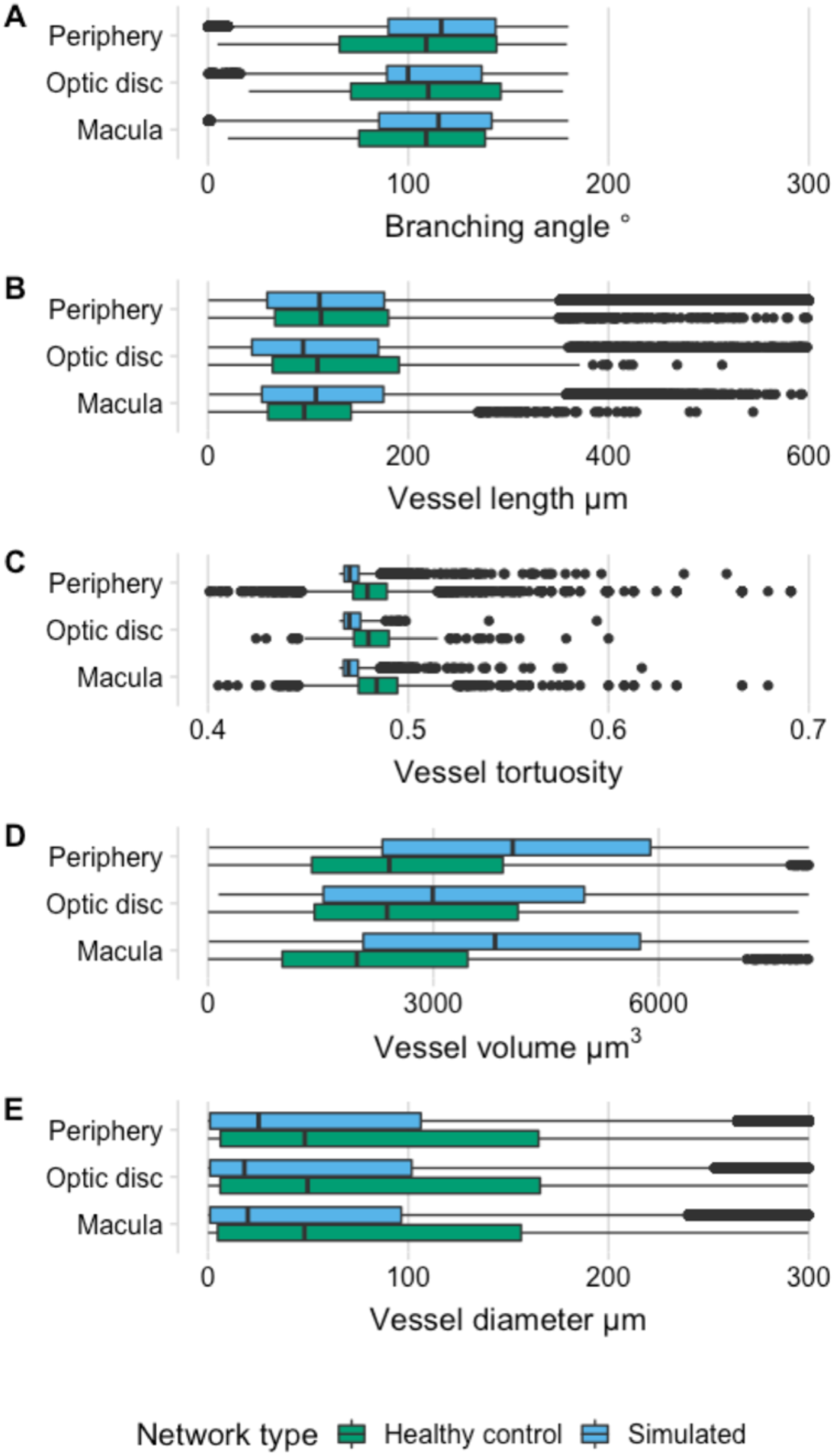
Comparison of retinal vascular geometry distributions visualised with bar plots between manually segmented networks from OCT-A data (normal volunteers not ascertained for disease status) and simulated networks. a) Branching angle, b) vessel length (µm), c) vessel tortuosity, d) vessel network volume and e) vessel diameter (µm), in three regions: macula (5.5mm diameter circular area centred on the fovea [33]), optic disc (3.6mm diameter area centred on the optic disc centre [34]), and periphery (all vessels outside those regions).

### Simulating retinal blood flow and validation using fluorescein angiography

We have previously developed a mathematical framework for simulating blood flow in three dimensional vascular networks, which uses one-dimensional Poiseuille flow [35]. Our simulated retinal networks are ideally suited for this framework, having just one arterial input and one venous outlet meaning that pressure boundary conditions can be easily specified. Setting arterial pressure by sampling from a normal distribution parameterised by mean = 56.2 mmHg, s.d. = 14.0 mmHg [36] and similarly for venous pressure with mean = 20.0 mmHg and s.d. = 10.0 mmHg [36] gave an average total retinal flow prediction of 34.4 ± 1.8 µL/min, which is slightly lower, but still in good agreement with reports in the literature from healthy retinas (for example, 45.6 ± 3.8 µL/min [36] 44 ± 13 µL/min [37] 50.7 µL/min [37] (**Figure 4a,b**).

**Figure 4:**
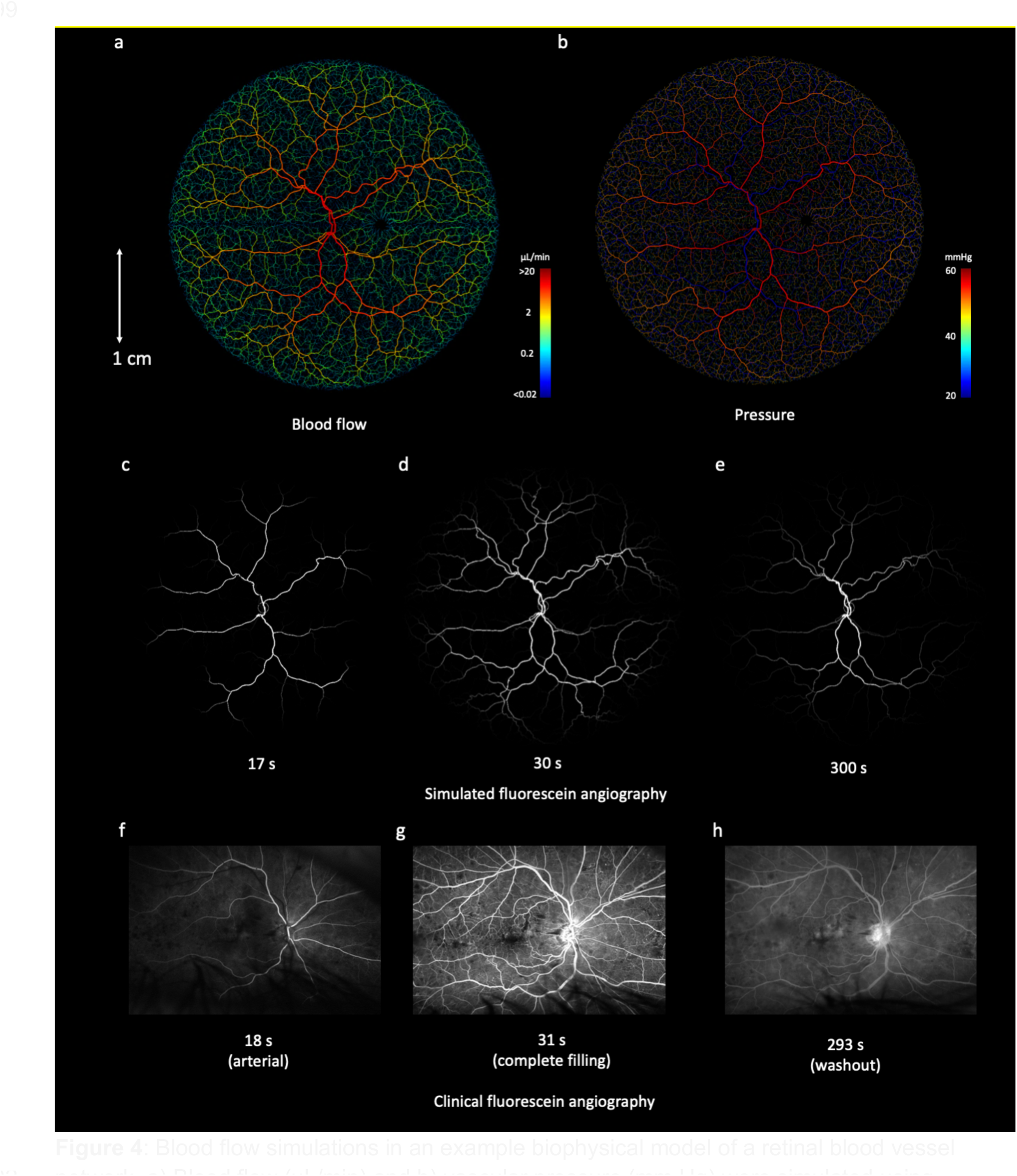
Blood flow simulations in an example biophysical model of a retinal blood vessel network. a) Blood flow (µL/min) and b) vascular pressure (mm Hg) were simulated using Poiseuille flow, with inlet arterial pressure and outlet venous pressure fixed at 56.2±14.0 and 20.0±10.0 mmHg, respectively. c-e) Simulated delivery of fluorescein at 17 s (arterial phase), 30 s (venous phase) and 600 s (recirculation), with clinical fluorescein images (registered to the same coordinate space) shown in f-h for comparison

To further evaluate these flow results, we next performed a simulation of retinal fluorescein delivery. Fluorescein angiography (FA) is used in ophthalmology for diagnosis of macular edema, macular degeneration, RVO, DR, and other diseases [38, 39]. Fluorescein is injected as a bolus into the median cubital vein, and 10-15 seconds later appears in the choroidal vasculature at the rear of the eye [40]. Within 2 seconds of this, fluorescein appears in the anterior arteries and arterioles, and a further two seconds later by partial filling of venules and veins, followed by total filling and recirculation.

We simulated the systemic pharmacokinetics of fluorescein using literature data (**Supplemental Figure 2**), with the passage of fluorescein modelled as two displaced Gaussian functions to model the first and second passes, and an exponential washout term corresponding to systemic extraction. This time course was propagated through our synthetic retinal networks by partitioning by flow at branch points and delaying according to cumulative velocities. The delay between arterial and venous filling with fluorescein, across 1000 simulation runs was 7.3 ± 0.7 s, which is in keeping with timings described in clinical data [40]. Visual inspection of fluorescein delivery also revealed a good accordance with clinical delivery profiles (**Figure 4c-h**).

### Simulating retinal pathology

Given the physiologically-realistic results provided by our flow models, we next sought to perturb our simulated networks to examine the effect of pathological changes. As a first demonstration, we simulated the effect of RVO. A random location of artery-vein crossover on a large retinal vein was reduced in diameter by 80%. Blood flow within the network was recalculated, revealing a large region of hypoperfusion, as expected. This strongly reflected the presentation of RVO found in clinical FA data (**Figure 5**) and induced a regional reduction in blood flow of 9.8 µL/min in the vessels immediately downstream of the occlusion.

**Figure 5:**
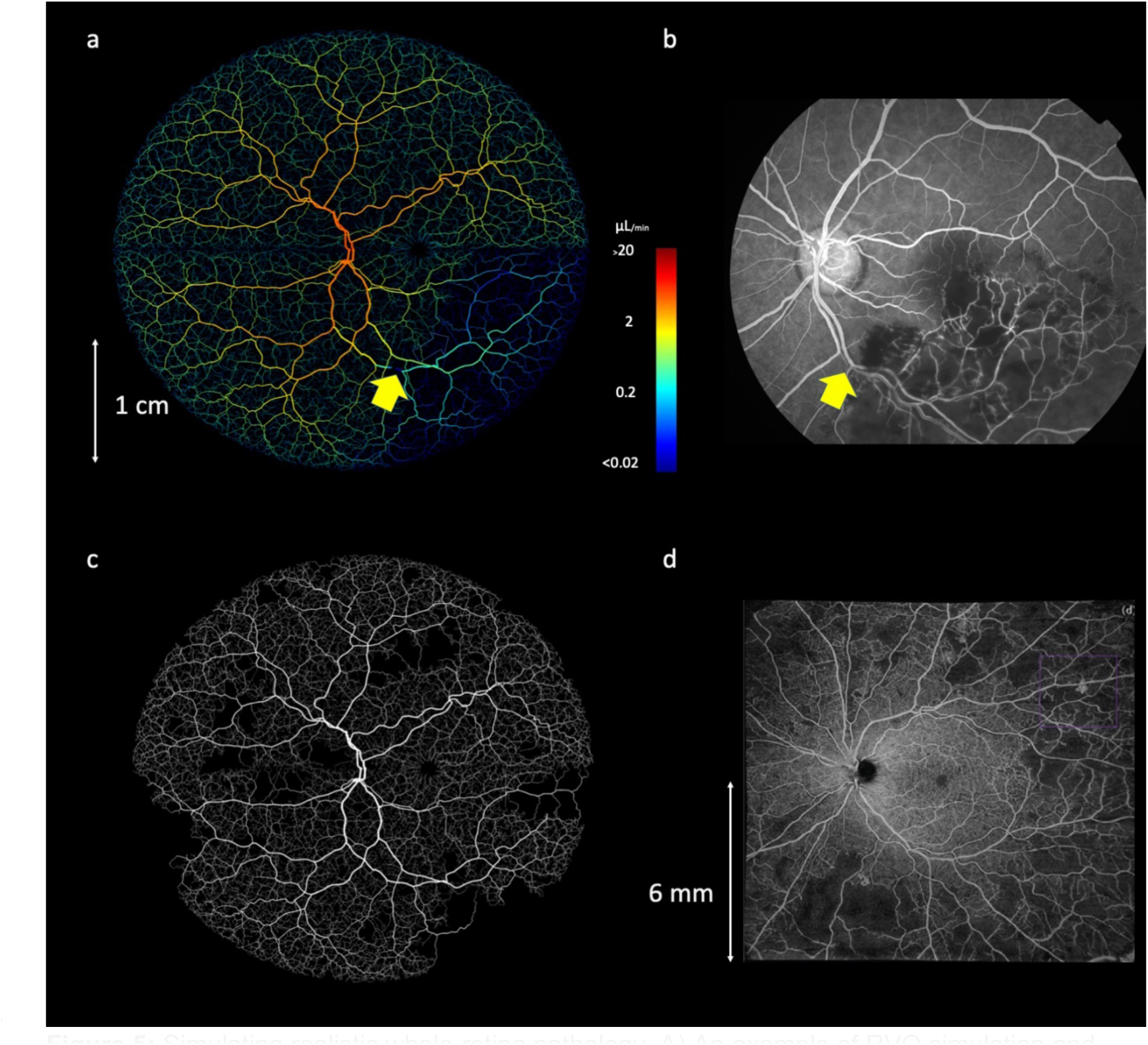
Simulating realistic whole-retina pathology. A) An example of RVO simulation and loss of flow in downstream vessels. The yellow arrow shows the location of an imposed 80% decrease in vein diameter. b) An FA image of retinal occlusion, revealing a similar pattern of perfusion loss as simulated in (a). c) The onset of DR, simulated by inhibiting flow in randomly-selected peripheral arterioles. D) An OCT-A image of a retina exhibiting stage 4 DR, evidenced by extensive loss of perfusion in vessels and regions of ischemia.

Next, we constructed a simple model of DR [41–43], in which arterioles with a radius less than 35 µm were randomly selected and occluded, and the resultant change in network flow calculated. All vessels that become non-perfused, either up- or downstream of the occluded vessel, were removed from the network, creating regions of ischemia, with occasional surviving vessels passing through (**Figure 5c-d, Supplemental Figure 3**). Occlusions were simulated in batches of 5, initially from the periphery (>1cm from the macula centre), and then at decreasing minimum distances from the macula, as typically found in the clinical presentation of DR.

Both our retinal occlusion model and DR model produced images that were highly reminiscent of clinical images of both pathologies (**Figure 5**), with loss of flow in downstream vessels in our RVO model and loss of perfusion and regions of ischaemia in the DR model.

### Generating synthetic clinical ophthalmology data with deep learning

Our next challenge was to use deep learning to define a mapping between our biophysical vascular model and clinical ophthalmology data (and vice versa). For this we used cycle-consistent generative adversarial networks that enabled the translation of image texture and style between image domains [44]. We undertook this for three clinical imaging modalities: OCT-A, retinal photographs and FA.

We embedded our synthetic retinas in three-dimensional grids with axial and lateral resolutions of 6.3 µm and 21 µm, respectively, to match our clinical OCT-A data. We then trained three cycle-consistent GANs on these synthetic retinas, with each GAN mapping the conversion between the synthetic images and a different imaging modality. 590 retinal photographs, 43 OCT-A en-face images and 570 FA images were used in training for this purpose. PI-GAN enabled the geometry of source images (simulations-domain A) to be translated into a target style (retinal photographs-domain B, OCT-A-domain C, and FA-domain D). As can be seen in **Figure 6a**, following 400 training epochs, the pattern of synthetic vasculature was realistically transferred into the style of each target image. The Frechet Inception Distance (FID) was 6.95 for retinal photographs, 5.17 for fluorescein angiographs and 3.06 for OCT-A en-face images, indicating a small distance between feature vectors for real and fake images.

**Figure 6:**
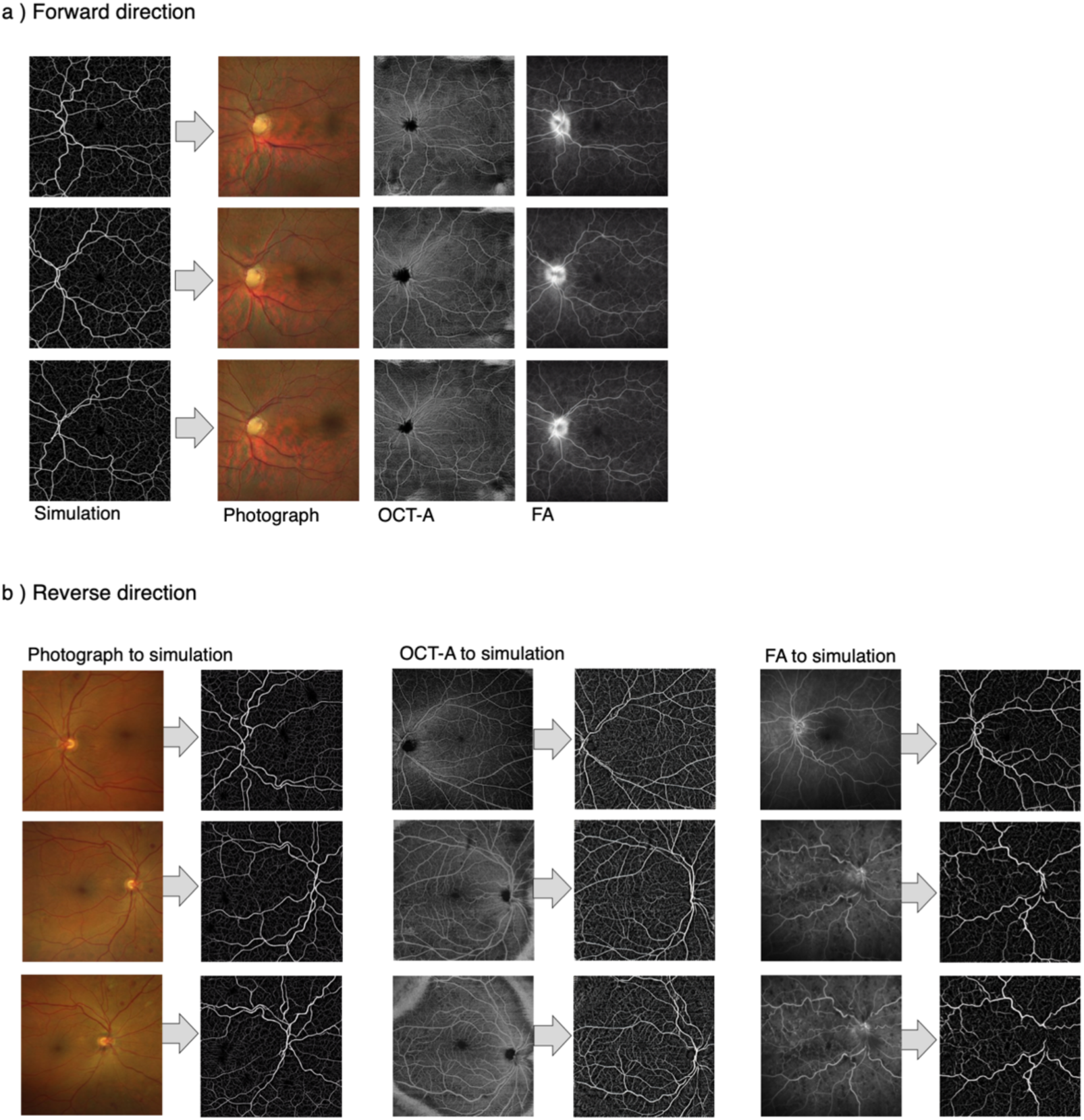
Generation of multi-modality retinal images from biophysical simulations, using physics-informed generative adversarial networks. Direction 1 involves conversion of domain A (simulated network) into domains B (fake retinal photograph), domain C (fake OCT-A) and domain D (fake fluorescein angiography). Direction 2 involves conversion of real retinal images (domains B-D) into fully connected networks/segmented data (domain A)

This process generated authentic-looking retinal image data with matched, fully specified ground truth blood vessels. However, cycle consistency also allows the reverse operation: to generate simulation data from clinical images (**Figure 6b**). This enabled blood vessel networks to be segmented from OCT-A images and compared with manual segmentations of the same data (**Figure 7**). Visual inspection of PI-GAN segmentations revealed many more small ‘elusive’ vessels [45] than represented within our manually-segmented images, arguably providing superior segmentation accuracy than the manual ‘gold standard’. Accordingly, the mean Dice score for OCT-A images was low (mean 0.35, s.d. 0.12 (2.s.f)), but the sensitivity (the percentage of pixels labelled as vessel in the manual segmentation that were also identified as vessel by PI-GAN) was high (87.1% (s.d. 1.20)), showing that PI-GAN is able to accurately label almost all of the vessels identified by human operators (**Figure 7g-j**).

**Figure 7:**
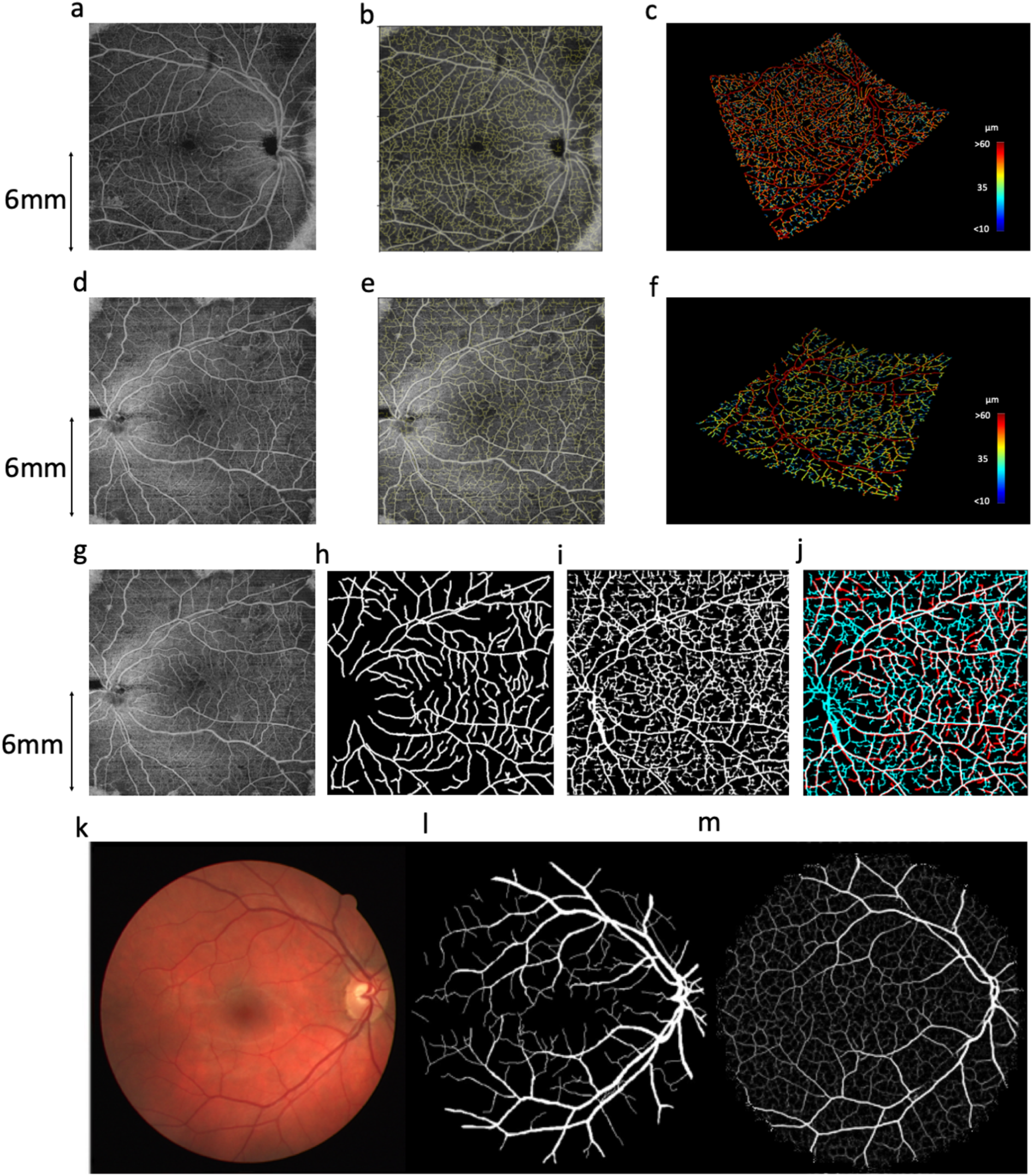
Blood vessel segmentation from OCT-A data with PI-GAN. A and d) OCT-A en-face images of retinal vasculature. B and e) The same OCT-A images with vessel segmentations from PI-GAN. C and f) Segmented vessels projected in three-dimensional space, colour-coded for vessel radius. G) An OCT-A image with h) manually-segmented and i) PI-GAN-segmented blood vessels. J) A composite image of manually- and PI-GAN-segmentations, with overlapping pixels rendered white, pixels with only PI-GAN-detected vessels in blue and pixels with only manual-detected vessels in red. K) A retinal photograph taken from the DRIVE data set [46] with l) manually-segmented and m) PI-GAN-segmented blood vessels.

To further investigate this result, we evaluated PI-GAN on two publicly available retinal photograph data sets with corresponding manual segmentations (STARE and DRIVE). Contrasting the widefield (130 degree and 200 degree montage) images analysed here, these datasets were acquired with a smaller 45 degree FOV, and are widely used in benchmarking vessel segmentation. Again, Dice scores comparing manual and PI-GAN segmentations were low but, as shown in **Figure 7k-m** and **Supplemental Figure 5**, PI-GAN was able to detect most of the manually-segmented vessels, but also many smaller, elusive vessels. Mean DICE score between DRIVE manually segmented data and segmentations generated using PI-GAN was 0.56 (s.d. 0.013) (2.d.p) and for STARE it was 0.64 (s.d. 0.19) (2.d.p). These results call into question how appropriate manual segmentation is as a gold standard in this setting, visual inspection suggests these additional small vessels are indeed physiological and were simply missed by manual segmentation.

These results demonstrate the key ability of physics-informed simulations with deep learning to autonomously segment blood vessels within a range of ophthalmology imaging modalities, without the need for any manually-labelled training data.

## Discussion

Methods that enable the quantitative assessment of retinal vasculature from clinical ophthalmology images are critically needed for evaluating the progression of diseases such as DR, and also to support research into the influence of systemic diseases such as cardiovascular disease and vascular dementia [3, 4]. In this regard, deep learning is rapidly transforming ophthalmology, but requires access to large volumes of well-curated data before it can be implemented with confidence in the clinic [47, 48]. In the assessment of vasculature, manual image labelling constitutes a considerable bottleneck in terms of time, expense and labelling accuracy [49], as manual segmentation of a single 2D retinal image can take multiple hours [50]. Inter and intra grader variability can also be significant within the segmentation process [51, 52]. Most segmentation studies have been conducted in 2D retinal fundus photographs using public datasets [12, 53, 54]. Approaches that can relieve this bottleneck are urgently needed to enable the robust translation of deep learning into the clinic.

To address these challenges, we have presented here a physics-informed, generative approach that combines biophysical simulation with deep generative learning. A useful outcome of this approach is the ability to automatically segment vascular data from clinical evaluation images, without any need for manual segmentation. Specifically, we created a linked set of algorithms that draw on established principles in biophysics to simulate fully-connected retinal vasculature, in a three-dimensional domain, with special treatment for optic disc and macular regions. The full connectivity of our models, with separate arterial and venous trees, enables realistic blood flow and delivery simulations (for example, as we show in fluorescein angiography). We demonstrated that our synthetic vascular networks are highly concordant with real retinal vasculature metrics, with network statistics matching those from manual segmentations, in three regions: the optic disc, macula and periphery.

This close accordance between simulation and real-world geometries is key to its ability to segment blood vessels from ophthalmology images. Cycle-consistent deep generative learning allowed us to create realistic fundus photograph, OCT-A and FA images that inherently maintained feature geometry through the translation from simulation to clinical image domains. The resultant data are inherently paired, and so could provide data to augment conventional supervised learning approaches. However, cycle-consistency also facilitates the reverse translation, from clinical image domains back into the simulation domain, allowing the automated segmentation of blood vessels without human-labelled data. Comparing segmentation performance against manual segmentations revealed a much greater ability to label small vessels, and with excellent overlap with larger manually-segmented vessels. However, overall performance assessed via DICE score showed a relatively low accordance, due in part to the greater ability of the PI-GAN approach to detect small blood vessels, but also false-positives in both human-labelled and PI-GAN-labelled vessels. In regard to the latter, there are cases where GANs can ‘hallucinate’ features in images [55].

To date, supervised deep learning approaches have yielded impressive results in 2D vessel segmentation relative to manual segmentation, although tend to favour precision over recall [5], resulting in an under-segmentation of faint vessels, underestimation of the width of thicker vessels and some ‘elusive’ vessels being missed [45]. This is problematic for diagnostic interpretation, because many biomarkers (such as artery-vein (AV) ratio, branching angles, number of bifurcations, fractal dimension and tortuosity) need precise measurements of individual vessels. GANs incorporating cycle-consistency have previously been used for medical imaging domain machine learning tasks such as chest MRI to X-ray CT transformation [16], PET image denoising [17], and artefact reduction in fundus photography [18]. Likewise Menten et al used the space colonisation algorithm to generate macular blood vessel images, which they coupled with deep learning [19].

Our approach builds on this by incorporating biophysically-informed models of flow within fully-connected artery and venous networks that extend across the entire retina, and our use of it to inform cycle-consistent deep generative learning. These developments allow application in larger field of view images (e.g. wide-field fundus photography), and also enable a large range of future applications, including flow modelling and oxygen delivery [56]. Moreover, given our ability to model arterial and venous trees, there is potential for independent segmentation of both vascular supplies.

These biophysical simulations also aimed to capture the wide range of variation found in real retinal networks, by varying the 26 simulation parameters across their reported physiological range. A further advantage of developing flow models into our biophysical framework was the ability to simulate pathology, such as the progression of DR and RVO. Many other pathologies could be simulated in follow-on studies, including changes in retinal vessel diameters associated with factors such as aging or hypertension. For example, Wong and colleagues reported retinal arteriolar diameters to decrease by approximately 2.1 µm for each decade increase in age, and by 4.4 µm for each 10 mmHg increase in arterial blood pressure [57]. Performing disease-specific deep generative learning runs will enable us to further refine our segmentation approaches and begin to characterise pathology.

Accordingly, there is also potential to use clinical data to further improve our biophysical simulations, enabling more accurate modelling of retinal physiology (and disease) and the ability to develop interpretable AI systems. The results of several recent studies using deep learning suggest that that retinal vasculature can provide a window into many systemic diseases (including dementia [3], kidney disease [58] and cardiovascular disease [4]), but cannot easily explain the structural basis of these associations. A PI-GAN framework is inherently coupled to biophysical laws, and so could help determine their origins or underpinning mechanistic processes. Additional challenges for segmentation are artery-vein classification [59] and establishing connectivity of the vessels [60], which, having a well-defined ground truth data set from simulations, could be realised through PI-GAN.

Overall our results demonstrate the potential of biophysical models of the retina, which can be interrogated to understand how physiological perturbations (such as disease) effect vascular function. Further work could explore regional variability in blood flow, with the temporal side exhibiting greater flow than the nasal side in both retinal venules and arteries, which may be related to retinal ganglion cell numbers [61]. Additionally, the model could be used in predicting inhibitors of angiogenesis, such as VEGF inhibitor Bevacizumab. Incorporating this model into a larger-scale retinal model (including the choroidal supply) would enable complete simulation of the retinal supply. The ability to then apply these simulation results for the interpretation of clinical images, via physics-informed generative learning, is a significant step forward.

## Acknowledgements

This research was funded by Cancer Research UK (C44767/A29458 and C23017/A27935) and EPSRC (EP/W007096/1). Audit number 1078 was used in accessing ophthalmological image data from Moorfields Eye Hospital NHS Foundation Trust. The audit was authorised by Moorfields Clinical Audit team.

The authors would like to thank Henry Cole, Andrew Kume, Jinyu Li, Kendra Hilliard, Yiyun Zhang, Jiahao Xu, Shuo Wu who assisted with data pre-processing, labelling and segmentation and the participants and patients who contributed ophthalmological imaging data.

## Author contributions

EEB contributed conception and design, analysis and interpretation of data, creation of software, and drafting the work. AG contributed creation of new software used. NH contributed analysis and interpretation of data. PS contributed creation of new software used. LG contributed analysis and interpretation of data. HC contributed analysis and interpretation of data. CW contributed analysis and interpretation of data. AM contributed creation of new software used. RS contributed conception and design, acquisition, and substantial revision to draft. RR contributed conception and design of work, acquisition of data, interpretation of data, and substantially revising draft. SWS contributed conception and design of work, acquisition, interpretation of data, creation of new software, and substantial revision of draft.

## Materials & correspondence

Professor Simon Walker-Samuel

## Code availability

Retina simulation software is available in https://github.com/CABI-SWS/vessel_sim, which has a dependency on https://github.com/AndrewAGuy/vascular-networks. Deep generative learning was performed using https://github.com/junyanz/pytorch-CycleGAN-and-pix2pix.

## Data availability

Simulated retinal networks are available in https://www.dropbox.com/scl/fo/whwru5rmz8g7cr0h8ytg1/h?rlkey=ynbh2kdhe0pcvpfo6cypm9oc6&dl=0.

## Methods

### Procedural generation of synthetic retinal vasculature

Generation of synthetic retinas followed multiple, length scale dependent steps. Firstly, the values of geometrical parameters were set by sampling from a normal or uniform distribution according to parameter values shown in **Supplemental Table 1.** Networks in the form of spatial graphs (i.e. branching nodes connected via vessel segments) were constructed using multiple, linked algorithms.

### Lindenmeyer system seeding

Firstly, seeding networks following approximate retinal vascular branching geometry were constructed, starting with a putative central retinal artery and retinal vein positioned at the centre of the optic disc. The diameters of the retinal artery and vein were 135 ± 15 μm and 151 ± 15 μm, respectively [23], oriented parallel to the optic nerve (defined as the z-direction). Two branches were added to the end of each of these segments, oriented in the x-y plane, and one directed above and the other below the retinal midline. Subsequent branching of these vessels was performed stochastically, with segment lengths between bifurcations set as a fixed fraction of vessel diameter (18 ± 3) and bifurcation vessel diameters set according to:

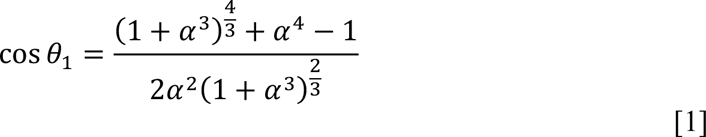

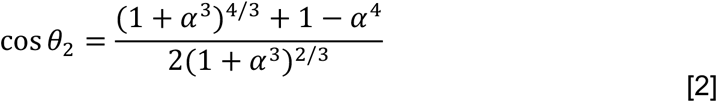

Normally-distributed noise was added to branching angle values, with a standard deviation of 5°. Vessel bifurcation angles were assigned such that the larger vessel oriented towards the macula to create putative major vessels oriented around the macula. Fifth-order bifurcations were added to the network, or until vessels breached the edge of the retina domain. **Supplemental Figure 1a** shows an example of an L-system seeding network.

### Major vessel growth

Seed vessel networks were used as input into a multi-scale growth algorithm for the creation of hierarchical vasculature. First, seed networks were amended to provide a uniform distribution of leaf nodes (terminating arteriole and venuole nodes created prior to construction of capillary networks at a later stage) throughout the circular domain, using Accelerated Constrained Constructive Optimisation [28], using a leaf node spacing of 3 mm. Multiscale, two-dimensional lattices were defined (stride lengths ranging from 3000 to 150 µm, with five iterations linearly spaced within that range) and used to grow vessel networks by progressively adding vessels into unoccupied lattice sites from neighbouring occupied sites, choosing the candidate vessel which minimised the expected change in network cost (see below), and progressively reducing the length scale when no more progress could be made. After the initial growth stage, all existing leaf nodes were removed [62]. At all stages of the major vessel growth the macula region was kept free of vessels by removing vessels which intersected it, forcing flow to divert around it.

As retinal vasculature is positioned in front of the retina itself, we optimised networks to minimise the area of the retina occluded by vessels, according to a cost function based on Murray’s law:

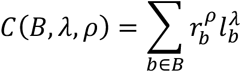

with ρ=1 and λ=1, and where *B* is the set of vessel segments in the network, with length *l* and radius *r*. After each growth step the network geometry was optimised by moving vessel nodes, and highly asymmetric bifurcations were trimmed for regrowth [28] using the thresholds from [27] to account for the high asymmetry of optimal networks [27]. After growth at each length scale was terminated, the networks were optimised topologically by allowing asymmetric bifurcations to move their low-flow side downstream and branches which were short compared to their expected length under the West, Brown and Enquist model [63] to be treated as a single higher-order split for regrouping using a method similar to [64]. Due to the two-dimensional nature of the networks, network self-intersections were tested for using the approach of [28] however, rather than resolving the intersections by making excursions around the contact site we rewire the vessels to prevent future iterations from recreating the same intersection.

Unlike the implementation of [28], leaf nodes were allowed to move from their nominal location up to a specified “pinning distance”, given as a fraction of the leaf spacing. Existing vessels could be specified as frozen, in which case the optimiser did not touch them. This approach was used to perturb the optimal root vessel structure with artificial tortuosity, strip away the downstream branches and regrow the downstream vessels, repeating this down the tree structure.

### Macula growth

Vessels supplying the macula have a characteristic radial structure, motivating the development of a particular approach to enforce this structure. This uses the same lattice site invasion approach between the macula outer radius and the fovea (which is kept vessel-free), but with the stride set low enough that the majority of the growth arises from spreading over many iterations at the same length scale rather than hierarchical refinement. The macula has a configurable flow rate density compared to the rest of the retina, ranging from 1.5 to 2.0 and leaf nodes are offset by uniformly sampling an offset in a disc around the nominal position to ensure that vessels did not align along the lattice sites. The macula vessels were prevented from doubling back on themselves by setting a hard limit on the vessel angle, preventing obviously non-physiological structures from arising whilst still allowing the radial pattern to develop. After all leaf nodes are created, a sparsity factor is specified and each leaf node removed with this probability, then the remaining vessels are geometrically optimised.

### Network overpass and interleaving

In the final stage, the arterial and venous networks have their collisions resolved using the method of [28], creating out-of-plane excursions around contact sites between the networks. To enable further micro-scale network growth techniques to create an interdigitated structure, we remove the low-flow side of all arterio-venous intersections with a radius below a critical value (5 um), leaving surviving vessel geometry untouched. Interdigitations were then created using a Space Colonisation implementation [65], interspersed with geometric optimisation.

### Vessel tortuosity

The multi-scale growth algorithm creates relatively straight paths between branching points, and to simulate tortuous retinal vessels, particularly in veins, sinusoidal displacements were overlaid. Two oscillations were superimposed according to:

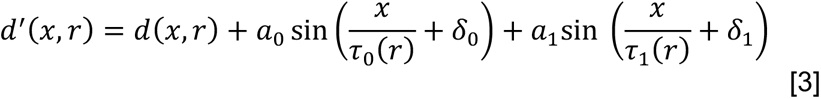

where *d*(*x,r*) is the path taken by a vessel with radius *r*, and *d*’(*x,r*) is the modulated path. The amplitude of displacements, a_0_+a_1_ ranged from *r* to 3.5*r* for arteries and *r* to 7.5*r* for veins, with a low frequency period (τ_0_, ranging from 15*r* to 25*r*) and a high frequency period (τ_1_, ranging from 30*r* to 50*r*). The phase of the modulations, *φ_0_* and *φ_1_,* enabled modulations to be matched between vessel bifurcations.

### Simulating vascular flow and fluorescein delivery

Blood flow in retinal networks were simulated using our REANIMATE platform [35], which uses a connectivity-based formalism to optimise Poiseuille flow in tree-like spatial graphs. As anterior retinal vasculature features a single arterial inlet and venous outlet, the system requires only one pressure boundary condition (the difference between arterial and venous inlet pressures), which was fixed at 56.2±14.0 and 20.0±10.0 mmHg, respectively.

Time-dependent delivery of contrast agent (e.g. fluorescein) was simulated as described in d’Esposito *et al* [35]. Briefly, a bolus of fluorescein was simulated according to

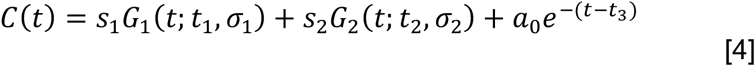

where *C*(*t*) is the concentration of fluorescein as a function of time *t*. The first two terms, Gaussian functions, represent the first and second pass of the bolus and the third term, an exponential decay, represents the washout phase [66] The width of the first and second pass were *σ*_1_ = 10 s and *σ*_2_ = 25 s, respectively, and the decay rate of the washout phase, β, was 0.043 /minute. *T*_1_, *t_2_* and *τ* are the time to peak for the first pass, second pass and washout phases, and were set at 0.171, 0.364 and 0.482 minutes, respectively [66]. *S*_1_, *s_2_* and α were fixed at 0.833, 0.336 and 1.064 (dimensionless units). Peak concentration was normalised to unity at the inlet to the retinal artery and the time course in each connected vessel segment was time-shifted according to the velocity of blood in each vessel and scaled according to the ratio of flow in the parent and child vessels at bifurcation points.

### Image datasets

This study was carried out in accordance with the Declaration of Helsinki [67]. Ethical approval of retrospective audit data was obtained through Moorfields Eye Hospital Research and Development Audit number 1078. Clinical ophthalmological retinal images were obtained from equipment at Moorfields Eye Hospital NHS Trust, London, UK: OCT-A images were obtained from a PLEX Elite 9000 (Carl Zeiss Meditec LLC, Dublin, CA, USA), ultra-wide true color retinal photographs were obtained from Zeiss Clarus 500 Fundus machine (Carl Zeiss Meditec LLC, Dublin, CA, USA), fluorescein angiograms were obtained from Optos widefield camera (Optos, Inc. Marlborough, MA, USA). 19 manually segmented OCT-A images were obtained from healthy controls not ascertained for disease status). These manual segmentations were used in comparison of network structure with simulated networks. Datasets of 570 FA images, 590 colour retinal photographs, 43 OCT-A en-face images, and 130 simulated networks were used in training and testing the PI-GAN algorithm.

### Manual labelling of clinical data

Manually labelled data was generated using a custom-built Python package enabling tracing of vasculature in 3D. The process involved placing user defining control points on the 2D image indicating where in a slab the vessel is located via maximum intensity projection. The z-height of the vessel was then fixed by identifying the height of the highest signal intensity voxel, which was manually constrained to exclude the choroid or RPE. The radius of each vessel was automatically calculated by setting a user-defined signal intensity threshold. Review of segmented structures was performed in 3D panel to assess and ensure labelling quality. In images with pathological blood vessels such as DR the abnormal vasculature or areas of neoangiogenesis were traced in the same manner. Vessel information (vessel coordinates, edge connectivity, number of edge points, edge point coordinates, radii, and vessel type) was exported and stored in Amira spatial graph format (ThermoFisher Scientific, Waltham, Massachussetts USA). Retinal regions were labelled. The macula was defined as a 5.5 mm diameter circular area centred on the fovea. The vessels surrounding the optic disc were labelled as a 3.6 mm diameter centred at the optic disc. Vessels outside these regions were defined as ‘peripheral’.

### Deep generative learning

Image-to-image translation was performed using cycle-consistent generative adversarial networks [18]. This algorithm enables automated unsupervised training with unpaired samples, learning a bi-directional mapping function between two different domains with deep generative adversarial networks. It utilises cycle consistency, where the reconstructed image obtained by a cycle adaptation is expected to be identical to the original image for both generative networks. Cycle-consistent GANs are composed of two main deep neural network blocks which are trained simultaneously: an image generator (generator) and an adversarial network (discriminator). There is a loss (G loss) to make a synthesised image from domain A closer to a real image from domain B, and a loss (D loss) to distinguish the synthesised image from domain A from a real image from A. There are also losses facilitating the conversion in the opposite direction (G loss making synthesised image from domain B closer to domain A, and D loss to distinguish synthesised and real domain B images. Additionally, cycle loss is the difference between the input image and the double-synthesised image and identity loss is the difference between output and input images. A train/validation/test split of 75%/5%/20% was used. All PI-GANtraining and evaluation was performed using a single NVIDIA Titan RTX GPU.

We iteratively trained a switchable PI-GAN algorithm with 500 epochs. All networks were trained using the optimizer ADAM solver [35] with β_1_ = 0.5, β_2_ = 0.999. The learning rate for the first 100 epochs was 2*10^−4^, and then linearly decayed to 2*10^−6^. Images were pre-processed with crop size 256 pixels. The minibatch size was 1. The loss weights λ were set as 10. The model was trained on NVIDIA TITAN RTX in Pytorch v1.9.1.

### Statistical evaluation of synthetic vessel networks

Vessel metrics of vessel branching angle, length, tortuosity, network volume and diameter were calculated. Analysis of variance (ANOVA) was used to assess differences in these metrics by retina region (optic disc, macula, and periphery) and by status (healthy control and simulated network) (Table 1) with eye (right OD/ left OS), participant sex, and scan pattern used as covariates.

### Evaluation metrics

#### Frechet inception distance

GAN output was evaluated using the Fréchet Inception Distance (FID), which evaluates model quality by calculating the distance between feature vectors for real and generated images. FID compares the distribution of generated images with distribution of real images that were used to train the generator. Lower FID scores indicate more similarity between two groups. The FID score is calculated by first loading a pre-trained Inception v3 model. The output layer of the model is removed and the output is taken as the activations from the last pooling layer, a global spatial pooling layer.

Three FID scores were calculated: real simulation images (domain A) versus manually segmented vasculature clinical images; real retinal photographs (domain B) versus PI-GAN generated retinal photographs; real OCT-A images (domain C) versus PI-GAN generated OCT-A images; real FA (domain D) versus PI-GAN generated OCT-A images.

#### Dice score

Dice scores were additionally calculated. This is a commonly used performance statistic for evaluating the similarity of two samples. For a ground truth segmentation label L and associated prediction P, we measure the binary Dice score D:

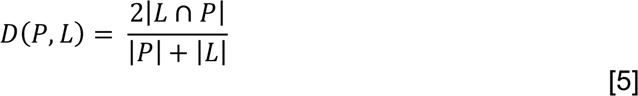

We carried out benchmarking of the PI-GAN algorithm against other models trained for manual segmentations from segmentation of retinal vessels using STARE, and DRIVE datasets public datasets, which are regularly used for benchmarking of algorithm results [46, 53]. DICE score were evaluated from the output of PI-GAN trained to carry out the mapping between simulated data segmentations and retinal photographs and compared to GAN performance without synthetic data.

## References

1. Shin, E.S., C.M. Sorenson, and N. Sheibani, Diabetes and retinal vascular dysfunction. J Ophthalmic Vis Res, 2014. 9(3): p. 362–73.

2. Trinh, M., M. Kalloniatis, and L. Nivison-Smith, Vascular Changes in Intermediate Age-Related Macular Degeneration Quantified Using Optical Coherence Tomography Angiography. Transl Vis Sci Technol, 2019. 8(4): p. 20.

3. Czako, C., et al., Retinal biomarkers for Alzheimer’s disease and vascular cognitive impairment and dementia (VCID): implication for early diagnosis and prognosis. Geroscience, 2020. 42(6): p. 1499–1525.

4. McClintic, B.R., et al., The relationship between retinal microvascular abnormalities and coronary heart disease: a review. Am J Med, 2010. 123(4): p. 374 e1–7.

5. Khanal, A., Estrada, R., Dynamic Deep Networks for Retinal Vessel Segmentation. Frontiers in Computer Science, 2020. 2.

6. Ting, D.S., G.C. Cheung, and T.Y. Wong, Diabetic retinopathy: global prevalence, major risk factors, screening practices and public health challenges: a review. Clin Exp Ophthalmol, 2016. 44(4): p. 260–77.

7. Aljuaid, A. and M. Anwar, Survey of Supervised Learning for Medical Image Processing. SN Comput Sci, 2022. 3(4): p. 292.

8. Ronneberger, O., Fischer, P., and Brox, T., U-net: convolutional networks for biomedical image segmentation. MICCAI (Freiburg im Breisgau), 2015.

9. De Fauw, J., et al., Clinically applicable deep learning for diagnosis and referral in retinal disease. Nat Med, 2018. 24(9): p. 1342–1350.

10. Hussain, S., et al., DilUnet: A U-net based architecture for blood vessels segmentation. Comput Methods Programs Biomed, 2022. 218: p. 106732.

11. Ren, K., et al., An improved U-net based retinal vessel image segmentation method. Heliyon, 2022. 8(10): p. e11187.

12. Jin, K., et al., FIVES: A Fundus Image Dataset for Artificial Intelligence based Vessel Segmentation. Sci Data, 2022. 9(1): p. 475.

13. de Moura, J., Novo, J., Ortega, M., & Charlón, P. 3D retinal vessel tree segmentation and reconstruction with OCT images. in Image Analysis and Recognition: 13^th^ International Conference, ICIAR 2016, in Memory of Mohamed Kamel. 2016. Póvoa de Varzim, Portugal: Springer International

14. Sherman, T.F., On connecting large vessels to small. The meaning of Murray’s law. J Gen Physiol, 1981. 78(4): p. 431–53.

15. Liu, Y., D. Zhang, and G.M. Karniadakis, Physics-informed generative adversarial networks for stochastic differential equations. SIAM Journal on Scientific Computing, 2020. 42(1): p. A292–A317.

16. Matsuo, H., et al., Unsupervised-learning-based method for chest MRI-CT transformation using structure constrained unsupervised generative attention networks. Sci Rep, 2022. 12(1): p. 11090.

17. Zhou, L., et al., Supervised learning with cyclegan for low-dose FDG PET image denoising. Med Image Anal, 2020. 65: p. 101770.

18. Yoo, T.K., J.Y. Choi, and H.K. Kim, CycleGAN-based deep learning technique for artifact reduction in fundus photography. Graefes Arch Clin Exp Ophthalmol, 2020. 258(8): p. 1631–1637.

19. Menten, M.J., Paetzold, J. C., Dima, A., Menze, B. H., Knier, B., & Rueckert, D. Physiology-based simulation of the retinal vasculature enables annotation-free segmentation of OCT angiographs. in Medical Image Computing and Computer Assisted Intervention–MICCAI 2022: 25th International Conference. 2022. Singapore: Springer Nature Switzerland.

20. Miller, D., Optics and Refraction : a User-Friendly Guide. 1991, Philadelphia, PA, USA: New York: Gower Medical Pub.

21. Mukherjee, P.K., Manual of Optics and Refraction. 2015, New Delhi: Jaypee Brothers Medical Publishers.

22. Lindenmayer, A., Mathematical models for cellular interactions in development. II. Simple and branching filaments with two-sided inputs. J Theor Biol, 1968. 18(3): p. 300–15.

23. Goldenberg, D., et al., Diameters of retinal blood vessels in a healthy cohort as measured by spectral domain optical coherence tomography. Retina, 2013. 33(9): p. 1888–94.

24. X. Liu, H.L., A. Hao, and Q. Zhao. Simulation of Blood Vessels for Surgery Simulators. In International Conference on Machine Vision and Human-machine Interface. 2010.

25. M. A. Galarreta-Valverde, M.M.G.M, C. Mekkaoui, and M. P. Jackowski. Three-dimensional synthetic blood vessel generation using stochastic L-systems. in Medical Imaging 2013: Image Processing, International Society for Optics and Photonics. 2013.

26. Buxbaum, W.S.a.P.F. Computer-optimization of vascular trees. in IEEE Transactions on Biomedical Engineering. 1993.

27. Schreiner, W., et al., The influence of optimization target selection on the structure of arterial tree models generated by constrained constructive optimization. J Gen Physiol, 1995. 106(4): p. 583–99.

28. Guy, A.A., et al., 3D Printable Vascular Networks Generated by Accelerated Constrained Constructive Optimization for Tissue Engineering. IEEE Trans Biomed Eng, 2020. 67(6): p. 1650–1663.

29. Luo, T., et al., Retinal Vascular Branching in Healthy and Diabetic Subjects. Invest Ophthalmol Vis Sci, 2017. 58(5): p. 2685–2694.

30. Smith, A.F., et al., Brain Capillary Networks Across Species: A few Simple Organizational Requirements Are Sufficient to Reproduce Both Structure and Function. Front Physiol, 2019. 10: p. 233.

31. Runions, A., Lane, B., Prusinkiewicz, P. Modelling Trees with a Space Colonization Algorithm in Eurographics Workshop on Natural Phenomena. 2007.

32. Tariq, A., A. Shaukat, and S.A. Khan. A Gaussian Mixture Model Based System for Detection of Macula in Fundus Images. in Neural Information Processing: 19th International Conference, ICONIP. 2012. Doha, Qatar: Springer Berlin Heidelberg.

33. Remington, L.A., & Goodwin, D., Clinical Anatomy and Physiology of the Visual System E-Book. Elsevier Health Sciences., 2021.

34. Quigley, H.A., et al., The size and shape of the optic disc in normal human eyes. Arch Ophthalmol, 1990. 108(1): p. 51–7.

35. d’Esposito, A., et al., Computational fluid dynamics with imaging of cleared tissue and of in vivo perfusion predicts drug uptake and treatment responses in tumours. Nat Biomed Eng, 2018. 2(10): p. 773–787.

36. Sun, R., et al., Central retinal artery pressure and carotid artery stenosis. Exp Ther Med, 2016. 11(3): p. 873–877.

37. Baumann, B., et al., Total retinal blood flow measurement with ultrahigh speed swept source/Fourier domain OCT. Biomed Opt Express, 2011. 2(6): p. 1539–52.

38. Savastano, M.C., et al., Fluorescein angiography versus optical coherence tomography angiography: FA vs OCTA Italian Study. Eur J Ophthalmol, 2021. 31(2): p. 514–520.

39. Marmor, M.F. and J.G. Ravin, Fluorescein angiography: insight and serendipity a half century ago. Arch Ophthalmol, 2011. 129(7): p. 943–8.

40. Ruia, S. and K. Tripathy, Fluorescein Angiography, in StatPearls. 2023: Treasure Island (FL).

41. Bek, T., Diameter Changes of Retinal Vessels in Diabetic Retinopathy. Curr Diab Rep, 2017. 17(10): p. 82.

42. Wang, W. and A.C.Y. Lo, Diabetic Retinopathy: Pathophysiology and Treatments. Int J Mol Sci, 2018. 19(6).

43. Tan, T.E., et al., Global Assessment of Retinal Arteriolar, Venular and Capillary Microcirculations Using Fundus Photographs and Optical Coherence Tomography Angiography in Diabetic Retinopathy. Sci Rep, 2019. 9(1): p. 11751.

44. Zhu, J.Y.P., T; Isola, P; Efros A.A., Unpaired image-to-image translation using cycle-consistent adversarial network. arXiv, 2017.

45. Zhou, Y., et al., A refined equilibrium generative adversarial network for retinal vessel segmentation. Neurocomputing, 2021. 437: p. 118–130.

46. DRIVE: Digital Retinal Images for Vessel Extraction.

47. Ting, D.S.W., et al., Artificial intelligence and deep learning in ophthalmology. Br J Ophthalmol, 2019. 103(2): p. 167–175.

48. Ting, D.S.W., et al., Deep learning in ophthalmology: The technical and clinical considerations. Prog Retin Eye Res, 2019. 72: p. 100759.

49. Day, T.G., J.M. Simpson, and R. Razavi, Improving image labelling quality. Nat Mach Intell, 2023. 5: p. 335–336.

50. Meng, X., et al., A framework for retinal vasculature segmentation based on matched filters. Biomed Eng Online, 2015. 14: p. 94.

51. Covert, E.C., et al., Intra- and inter-operator variability in MRI-based manual segmentation of HCC lesions and its impact on dosimetry. EJNMMI Phys, 2022. 9(1): p. 90.

52. Veiga-Canuto, D., et al., Comparative Multicentric Evaluation of Inter-Observer Variability in Manual and Automatic Segmentation of Neuroblastic Tumors in Magnetic Resonance Images. Cancers (Basel), 2022. 14(15).

53. Hoover, A., V. Kouznetsova, and M. Goldbaum, Locating blood vessels in retinal images by piecewise threshold probing of a matched filter response. IEEE Trans Med Imaging, 2000. 19(3): p. 203–10.

54. Staal, J., et al., Ridge-based vessel segmentation in color images of the retina. IEEE Trans Med Imaging, 2004. 23(4): p. 501–9.

55. Cohen, J.P., Luck, M., Honari, S. Distribution Matching Losses Can Hallucinate Features in Medical Image Translation. in Medical Image Computing and Computer Assisted Intervention – MICCAI 2018. 2018. Springer.

56. Sweeney, P.W., S. Walker-Samuel, and R.J. Shipley, Insights into cerebral haemodynamics and oxygenation utilising in vivo mural cell imaging and mathematical modelling. Sci Rep, 2018. 8(1): p. 1373.

57. Wong, T.Y., et al., Retinal vessel diameters and their associations with age and blood pressure. Invest Ophthalmol Vis Sci, 2003. 44(11): p. 4644–50.

58. Yeung, L., et al., Early retinal microvascular abnormalities in patients with chronic kidney disease. Microcirculation, 2019. 26(7): p. e12555.

59. Galdran, A., et al., State-of-the-art retinal vessel segmentation with minimalistic models. Sci Rep, 2022. 12(1): p. 6174.

60. Shin, S.Y., et al., Deep vessel segmentation by learning graphical connectivity. Med Image Anal, 2019. 58: p. 101556.

61. Garhofer, G., et al., Retinal blood flow in healthy young subjects. Invest Ophthalmol Vis Sci, 2012. 53(2): p. 698–703.

62. Schwen, L.O. and T. Preusser, Analysis and algorithmic generation of hepatic vascular systems. Int J Hepatol, 2012. 2012: p. 357687.

63. Brown, J.H., G.B. West, and B.J. Enquist, Yes, West, Brown and Enquist’s model of allometric scaling is both mathematically correct and biologically relevant. Functional Ecology, 2005.

64. Georg, M., T. Preusser, and H.K. Hahn, Global Constructive Optimization of Vascular Systems’, All Computer Science and Engineering Research. All Computer Science and Engineering Research, 2010. WUCSE-2010-11: p. 16.

65. Runions, A., Lane, B., & Prusinkiewicz, P. Modeling Trees with a Space Colonization Algorithm. 2007.

66. Parker, G.J., et al., Experimentally-derived functional form for a population-averaged high-temporal-resolution arterial input function for dynamic contrast-enhanced MRI. Magn Reson Med, 2006. 56(5): p. 993–1000.

67. General Assembly of the World Medical, A., World Medical Association Declaration of *Helsinki*: ethical principles for medical research involving human subjects. J Am Coll Dent, 2014. 81(3): p. 14–8.

