## Supplemental material for "Physics-informed deep generative learning for quantitative assessment of the retina"

**Supplementary material**

| **Parameter** | **Value** |
|---|---|
| *Eye geometry* | |
| Optic cup diameter ($O_d$) | 0.7 – 1.2 mm |
| Optic disc radius | (1.1 – 1.5) $O_d$ |
| Eye radius | 23 — 25 mm |
| Optic nerve – fundus displacement | 3.5 – 5.5 mm [72] |
| Fovea radius | 500 ± 20 μm |
| Macula radius | 2500 ± 200 μm |
| Retina radius | 30 – 40 mm [73] |
| *L-system* | |
| Central retinal artery diameter | 135 ± 15 μm |
| Central retinal vein diameter | 151 ± 15 μm |
| Central retinal artery initial branching angle | 10 ± 5 ° |
| Length of first retinal artery / vein segment | 0 — 1500 μm |
| L-system branching angle | 30 ± 2 ° |
| L-system inter-branch distance | (30 ± 3) $r_i$ |
| Optic disc / fovea angle | 6.3 ± 3.0 ° [74] |
| *CCO / LVM* | |
| Murray exponent | 2.4 ± 0.1 |
| Macula flow factor | 1.5 — 2 |
| Maximum spacing | 2500 μm |
| Minimum spacing | 150 μm |
| Number of refinements | 5 |
| *Tortuosity:* $a_1 \sin(x/p_1) + a_2 \sin(x/p_2)$ | |
| $p_1$ | (15 – 25) $r_i$ |
| $p_2$ | (30 – 50) $r_i$ |
| $a_1$ (artery) | (1.0 – 3.5) $r_i$ |
| $a_1$ (vein) | (1.0 – 7.5) $r_i$ |
| $a_2$ | $a_1$ * 0.4 |
| *Fluid dynamics* | |
| Central retinal artery pressure | 60 ± 5 mmHg |
| Central retinal vein pressure | 20 ± 5 mmHg |

**Supplemental Table 1.** Parameters used in our procedural modelling of retinal vasculature informed by metrics from clinical studies [75, 76]. Parameters expressed as ranges were treated as uniform random distributions, whereas values with uncertainties were treated as random normal distributions (mean ± s.d.). $r_i$ is the radius of the $i^{th}$ vessel segment; $x$ is distance along the vessel segment.

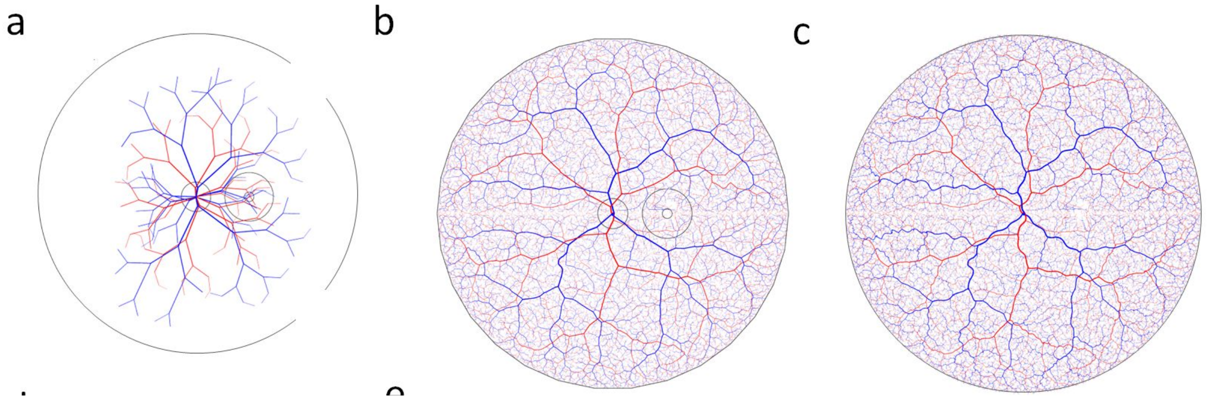


**Supplemental Figure 1**: Generation of synthetic retinal vascular networks. A) An example seeding L-system output, featuring prototype arterial (red) and venous (blue) trees emanating from a central optic disc, and with asymmetric branching towards a macular region offset to the right of the optic disc. B) A result of CCO and LSV algorithms applied to the networks in a). c) Overlay of tortuosity displacements using sinusoidal harmonics on (b). d) Three- dimensional projection of the simulation onto a model of retinal layers, featuring optic disc and macular structures modelled with split Gaussians.

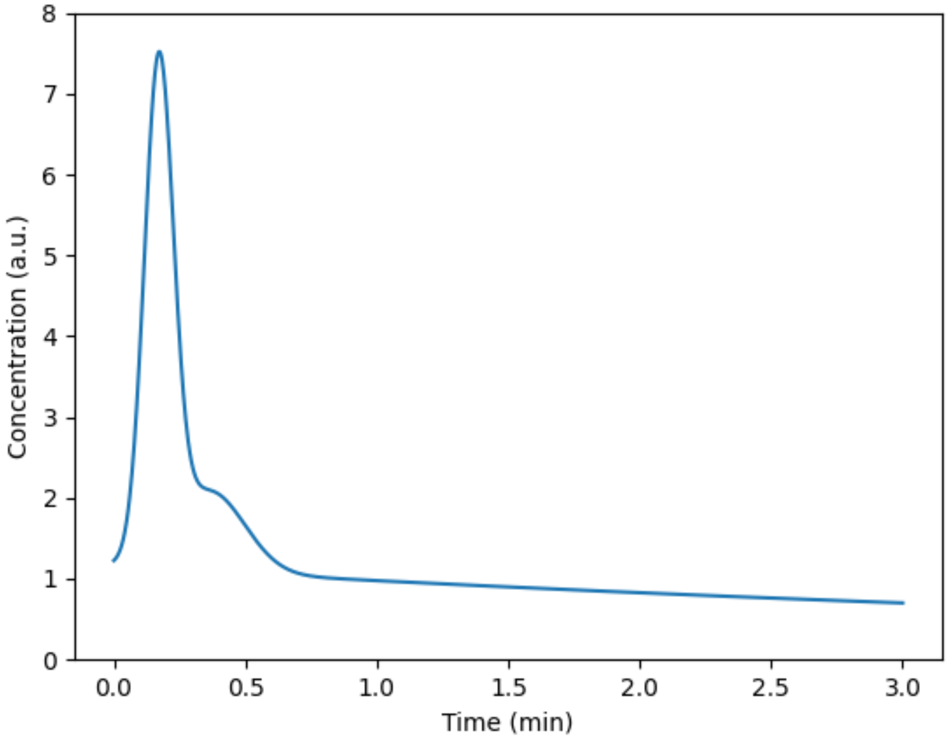


**Supplemental Figure 2.** Fluorescein arterial input pharmacokinetics used to simulate retinal fluorescein delivery.

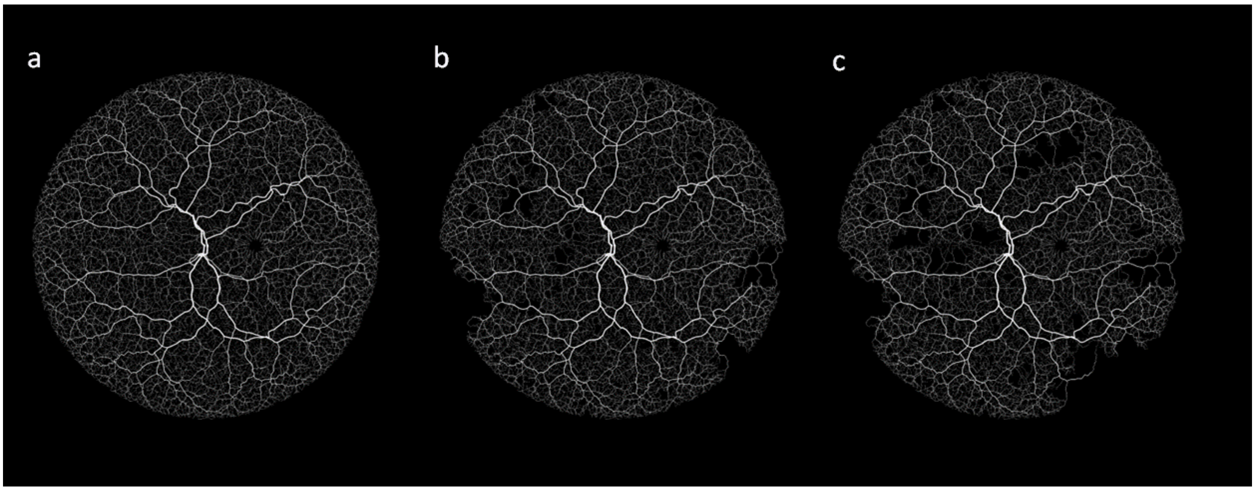


**Supplemental Figure 3.** Simulated progressive onset of DR in simulated retinal blood vessel networks.

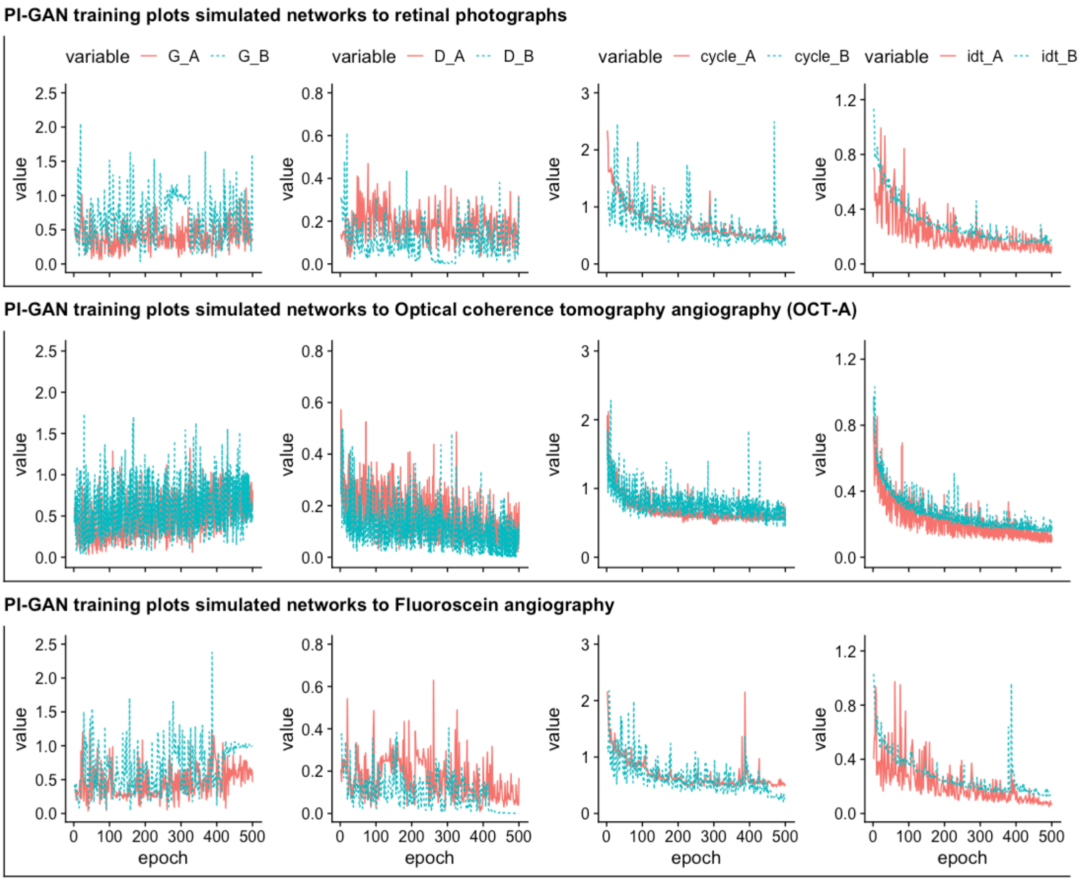


**Supplemental Figure 4**. PI-GAN training plots showing generator losses (G_A, generator loss for conversion of images from domain A to domain B (A→B) and G_B, generator loss for conversion of images from domain B to domain A (B→A)), discriminator losses (D_A, discriminator loss (G_A(A), D_B discriminator loss G_B(B)), cycle consistency (cycle_A (equation lambda_A * ||G_B(G_A(A)) - A||) and cycle_B (equation lambda_B * ||G_A(G_B(B)) - B||) and identity losses defined by equation lambda_identity * (||G_A(B) - B||

* lambda_B + ||G_B(A) - A|| * lambda_A) (idt_A, idt_B). Plots show the losses during training of simulated networks to retinal photographs, OCT-A images, and fluorescein angiography.

| | Mean (standard deviation) | | | | | | Statistical analys (p values) |
|---|---|---|---|---|---|---|---|
| | Periphery | | Optic disc | | Macula | | Real v simulated |
| | Healthy control (n=19) | Simulated (n=100) | Healthy control (n=19) | Simulated (n=100) | Healthy control (n=19) | Simulated (n=100) | ANOVA p value |
| Branching angle (degrees) | 104 (45.30) | 114 (40.0) | 107 (42.8) | 106 (39.4) | 106 (39.7) | 110 (42.1) | 0.82 |
| Inter-branch length (μm) | 136 (95.6) | 128 (92.3) | 143 (146.0) | 151 (182.0) | 112 (88.5) | 125 (75.6) | 0.17 |
| Tortuosity | 0.485 (0.0437) | 0.557 (0.325) | 0.488 (0.0517) | 0.552 (0.314) | 0.488 (0.0345) | 0.480 (0.0345) | 0.095 |
| Network volume ( $\mu m^3$) | 37058 (48860) | 34235 (127146) | 36359 (41876) | 52145 (182394) | 34603 (45212) | 34243 (127251) | 0.061 |
| Diameter ( μm) | 100.68 (179.50) | 92.11 (150.49) | 111.76 (235.23) | 101.58 (198.78) | 87.43 (296.70) | 62.42 (220.18) | 0.59 |

**Supplemental table 2**. Summary statistics (mean, standard deviation) for retinal vessel branching angle, inter-branch length, tortuosity, volume and diameter in three regions of the retina (periphery, optic disc, macula) and by data type (control, DR, simulation) and ANOVA p values.

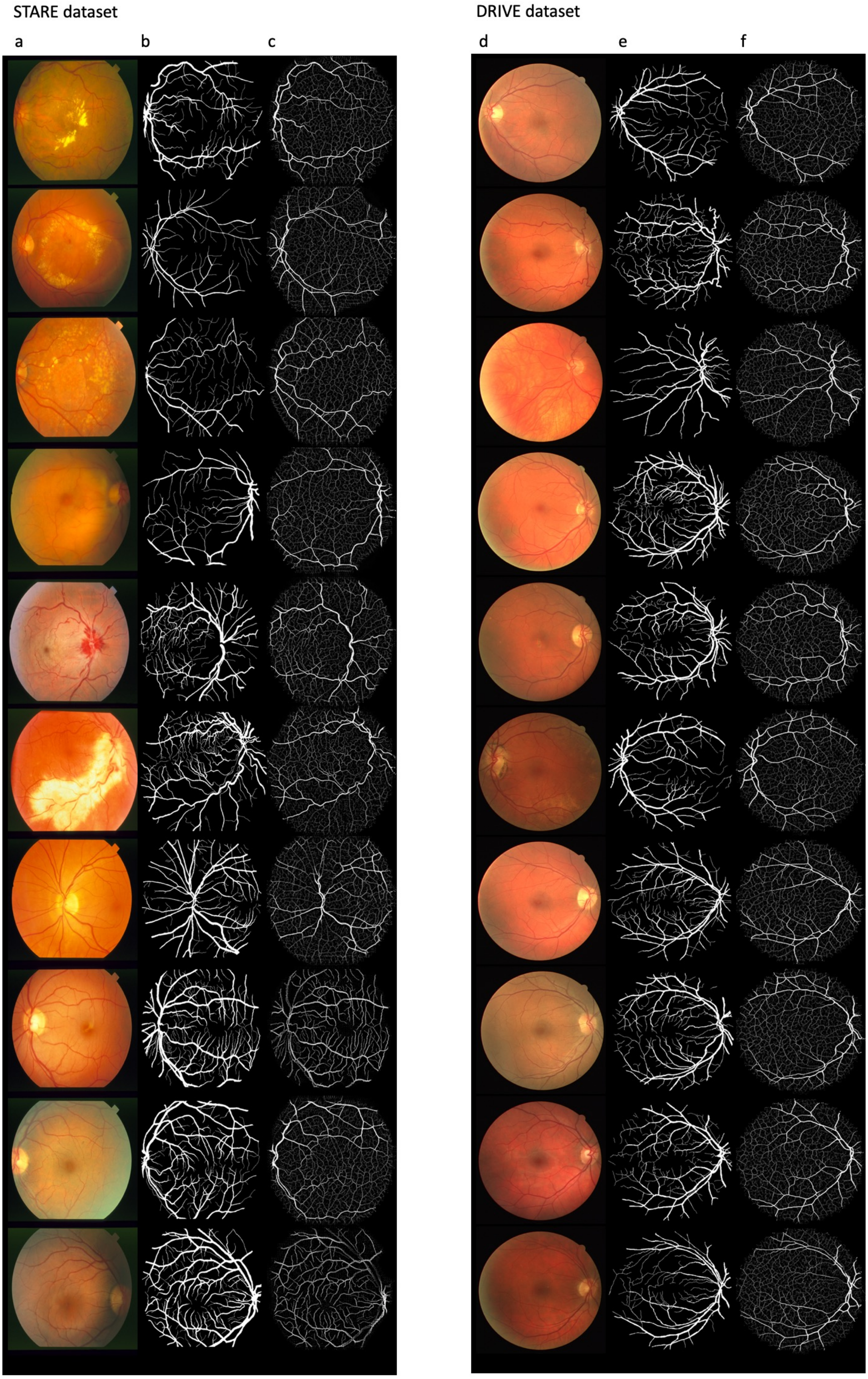


**Supplemental Figure 5.** Segmentation output of STARE and DRIVE images. a and d) original fundus images from STARE and DRIVE datasets respectively. b and e) manual segmentation image from the public datasets. c and f) PI-GAN based segmentation output.

**Supplemental video 1.** Video visualisation showing synthetic retinal artery vein vasculature and the simulated pharmacokinetics of fluorescein delivery
